# Two essential replicative DNA polymerases exchange dynamically during DNA replication and after replication fork arrest

**DOI:** 10.1101/364695

**Authors:** Yilai Li, Ziyuan Chen, Lindsay A. Matthews, Lyle A. Simmons, Julie S. Biteen

## Abstract

The replisome is the multi-protein complex responsible for faithful replication of chromosomal DNA. Using single-molecule super-resolution imaging, we characterized the dynamics of three replisomal proteins in live *Bacillus subtilis* cells: the two replicative DNA polymerases, PolC and DnaE, and a processivity clamp loader subunit, DnaX. We quantified the protein mobility and dwell times during normal replication and following both damage-independent and damage-dependent replication fork stress. With these results, we report the dynamic and cooperative process of DNA replication based on changes in the measured diffusion coefficients and dwell times. These experiments show that the replisomal proteins are all highly dynamic and that the exchange rate depends on whether DNA synthesis is active or arrested. Our results also suggest coupling between PolC and DnaX in the DNA replication process, and indicate that DnaX provides an important role in synthesis during repair. Furthermore, our results show that DnaE provides a limited contribution to chromosomal replication and repair.

## Introduction

The process of DNA replication is essential for all organisms. Failure to replicate DNA accurately in multicellular systems can lead to disease states or even cell death, while in bacteria, problems in DNA replication can affect cell fitness and viability (1,2). The replisome is a multiprotein complex that functions to replicate chromosomal and plasmid DNA; in bacteria, we define the replisome as a molecular machine that includes the DNA polymerase(s), the *β*-sliding clamp, the *β*-clamp loader complex, the helicase, the primase, and the single-stranded DNA binding protein (SSB) (3). Overall, the bacterial replisome has been investigated at length *in vitro*, yet few studies have used single-molecule approaches to understand how the replisome functions *in vivo* (3-8).

The replisome of *Bacillus subtilis*, a model organism for Gram-positive bacteria, has been particularly well studied (3,7,8). Unlike the replisome in Gram-negative bacteria including *Escherichia coli*, the *B. subtilis* replisome is confined to a specific position in the cell (9,10), and the DNA templates are pulled into the replisome during synthesis (8,11,12). Moreover, the *B. subtilis* replisome requires two different DNA polymerases, PolC and DnaE, for replication (13), which is more similar than the replication system of *E. coli* to the two-polymerase mechanism of eukaryotes (14,15). *In vitro* experiments on the *B. subtilis* replisome suggest that PolC is responsible for all leading DNA strand synthesis and most of the lagging DNA strand synthesis, whereas DnaE, an error-prone DNA polymerase (16,17), extends the lagging strand RNA primer before handing off to PolC (3). However, the *in vitro* model of DNA replication is incomplete, and high-resolution studies in bacteria are furthering our understanding of how DNA replication functions in the context of a living cell. For instance, Seco et al. recently reported that DnaE might be involved in *B. subtilis* leading strand synthesis (18), while Paschalis et al. proposed that DnaE contributes substantially to elongation during chromosomal replication (19). Moreover, other replisomal proteins including SSB and the *β*-sliding clamp can modulate the activity and fidelity of DnaE polymerase, suggesting that DnaE is capable of replicating substantial amounts DNA in live *B. subtilis* cells (19). Therefore, the contribution of DnaE to chromosomal replication remains unclear *in vivo*.

Despite many *in vitro* and *in vivo* studies of the *B. subtilis* replisome, the architecture and dynamical interactions of the replisome components in live *B. subtilis* cells remain unclear, and the functions during the normal DNA replication process of some key replisomal proteins including DnaE are still unresolved. Recently, the development of single-molecule imaging has provided a new and important tool to study DNA replication in living cells with high sensitivity and spatial resolution (18,20-23). In our previous work, we characterized the stoichiometry and the dynamics of one of the major DNA polymerases, PolC, in live *B. subtilis* cells (23). This work showed that PolC is highly dynamic, moving to and from the replication fork with three molecules present at each fork. Because replisome proteins work cooperatively to replicate DNA, it is unclear if the single-molecule behavior we observed for PolC is representative of other replisomal proteins. Furthermore, if DnaE has prominent roles in chromosomal replication or repair, a high-resolution approach should be capable of detecting such activity. Thus, more contextual information is needed to understand the *in vivo* architecture and behavior of the DNA replication machine in *B. subtilis*.

Here, we use super-resolution microscopy and single-particle tracking to determine the localization and dynamics of three replisomal proteins during normal DNA replication: the PolC and DnaE replicative DNA polymerases and the DnaX subunit of the *β*-sliding clamp loader complex. Based on these measurements, we elucidate the DNA replication mechanism by examining the localization and dynamics of the replisomal proteins during normal DNA replication and when the DNA replication is arrested via two distinct mechanisms: PolC disruption with 6-hydroxy-phenylazo-uracil (HPUra) (24) or cross-linking with mitomycin C (MMC) (25). Overall, the subcellular positioning, motion, and responses to DNA replication arrest indicate that the replisomal proteins exchange dynamically during DNA replication and that protein exchange dynamics change when DNA replication is arrested. Our results provide new insight into the molecular exchange of replisomal proteins during active DNA synthesis and after both damage-dependent and damage-independent fork arrest.

## Results

### Dynamical positioning of replisome proteins in live *B. subtilis* cells during normal DNA replication

To study the localization and dynamics of the replisome proteins, we genetically engineered fusions of the photoactivatable red fluorescent protein PAmCherry to the C-terminus of PolC (JWS213), DnaE (LAM380.1), or DnaX (LYL001) as the sole source of these essential proteins at the native locus of their respective genes. The PolC-PAmCherry strain also includes an ectopically expressed DnaX-mCitrine fusion under control of a xylose promoter, as a marker of the replisome position (20). We stochastically photoactivated 1 – 3 PAmCherry molecules per cell at a time, imaged this photoactivated subset until all PAmCherry was photobleached, then photoactivated a new subset. Photoactivated localization microscopy (PALM) super-resolution images were constructed after 5 – 10 iterations of this photoactivation-imaging-photobleaching cycle. For the PolC-PAmCherry strain, DnaX-mCitrine foci were imaged before any PAmCherry photoactivation. Consistent with earlier findings (20), DnaX-mCitrine forms clusters at the mid-cell or quarter-cell positions (Fig. 1A). PolC-PAmCherry molecules are enriched around the replisome area, although a significant number of molecules are distributed distal to the replisome throughout the cell (Fig. 1A).

**Figure 1.**
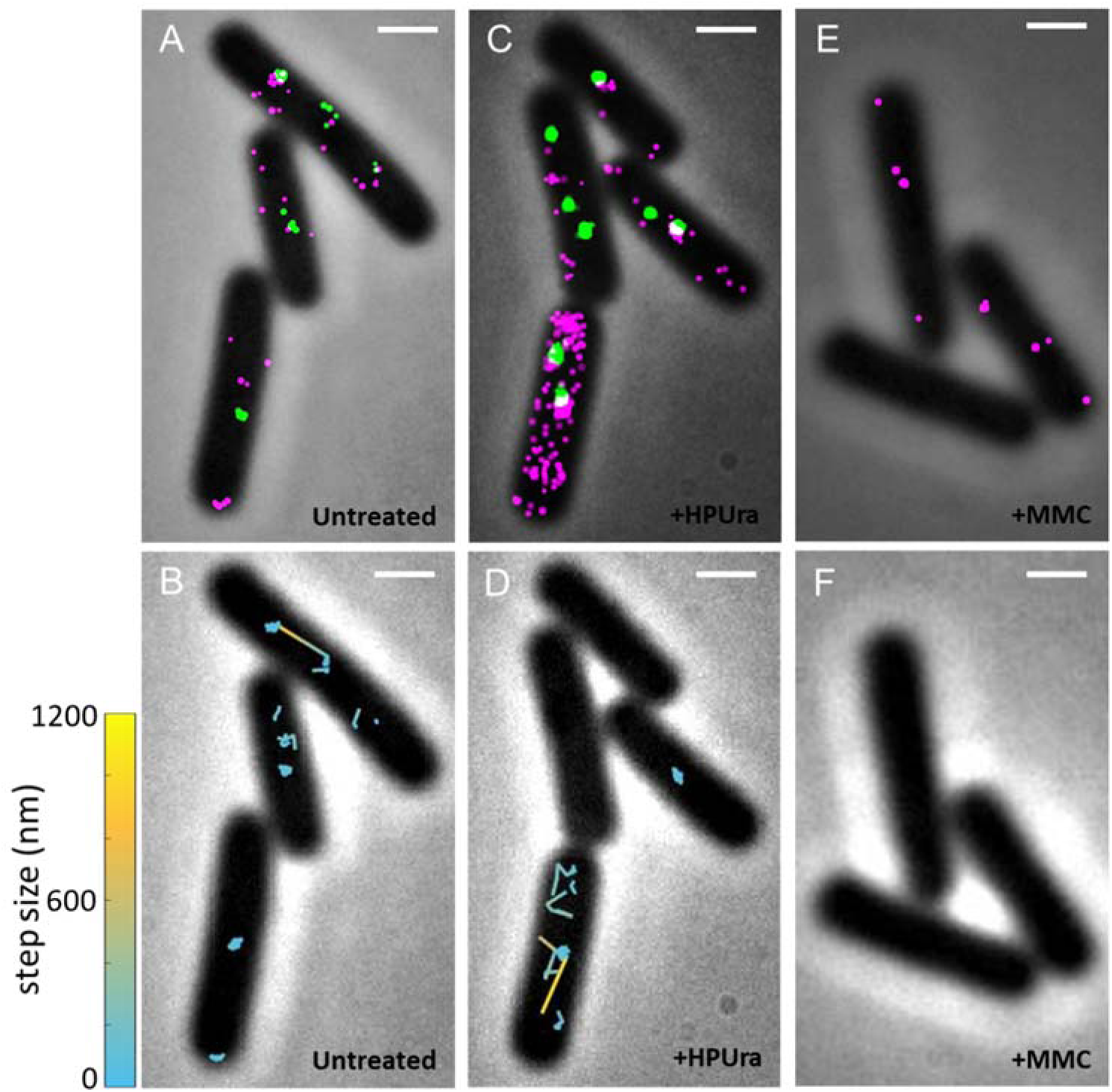
Location and dynamics of PolC in representative live *B. subtilis* cells. (A) Photoactivated localization microscopy (PALM) reconstructions of PolC-PAmCherry (magenta) overlaid with DnaX-mCitrine clusters (green) in untreated cells. (B) Representative trajectories of PolC-PAmCherry in untreated cells. The trajectories are color-coded by step size to distinguish the fastest motion (yellow) from the slowest motion (cyan). (C) PALM reconstructions of PolC-PAmCherry (magenta) and DnaX-mCitrine (green) in HPUra-treated cells. (D) Representative trajectories of PolC-PAmCherry in HPUra-treated cells color-coded by step size. (E) PALM reconstructions of PolC-PAmCherry (magenta) in MMC-treated cells. (F) PolC-PAmCherry molecules moved too fast to be tracked in MMC-treated cells. Scale bar: 1 μm.

We measured the real-time dynamics of PolC-PAmCherry molecules in these living cells using single-particle tracking (Fig. 1B, Movie S1). This mobility was very heterogeneous, thus we differentiated between two sorts of dynamical behaviors—one fast and one slow—by fitting the cumulative probability distribution (CPD) of the squared step sizes to a diffusion model with two mobile terms (Eq. 2, Methods). This analysis provided the average fast and slow diffusion coefficients, *D*_*PolC*-*fast*_ and *D*_*PolC*-*slow*_, respectively, corresponding the two PolC subpopulations. By analyzing 1230 tracks (SI Fig. S1, Fig. 2A), we found that although about 73% of PolC molecules move slowly in the cells (*D*_*PolC*-*slow*_ = 0.026 ± 0.005 μm^2^/s), a significant amount (27%) of PolC molecules diffuse much more rapidly, with *D*_*PolC*-*fast*_ = 0.5 ± 0.2 μm^2^/s (Table 1). To understand how the motion varies with position within the cell, we mapped the step sizes of PolC as a function of distance from the replisome (SI Fig. S1). We found that the step sizes of PolC decrease near the DnaX foci, implying that the slowly diffusing PolC-PAmCherry molecules correspond to PolC actively engaged in replication or at least co-localized to the replisome, whereas the fast population corresponds to PolC transiently coming in contact with chromosomal DNA away from the replisome.

**Figure 2.**
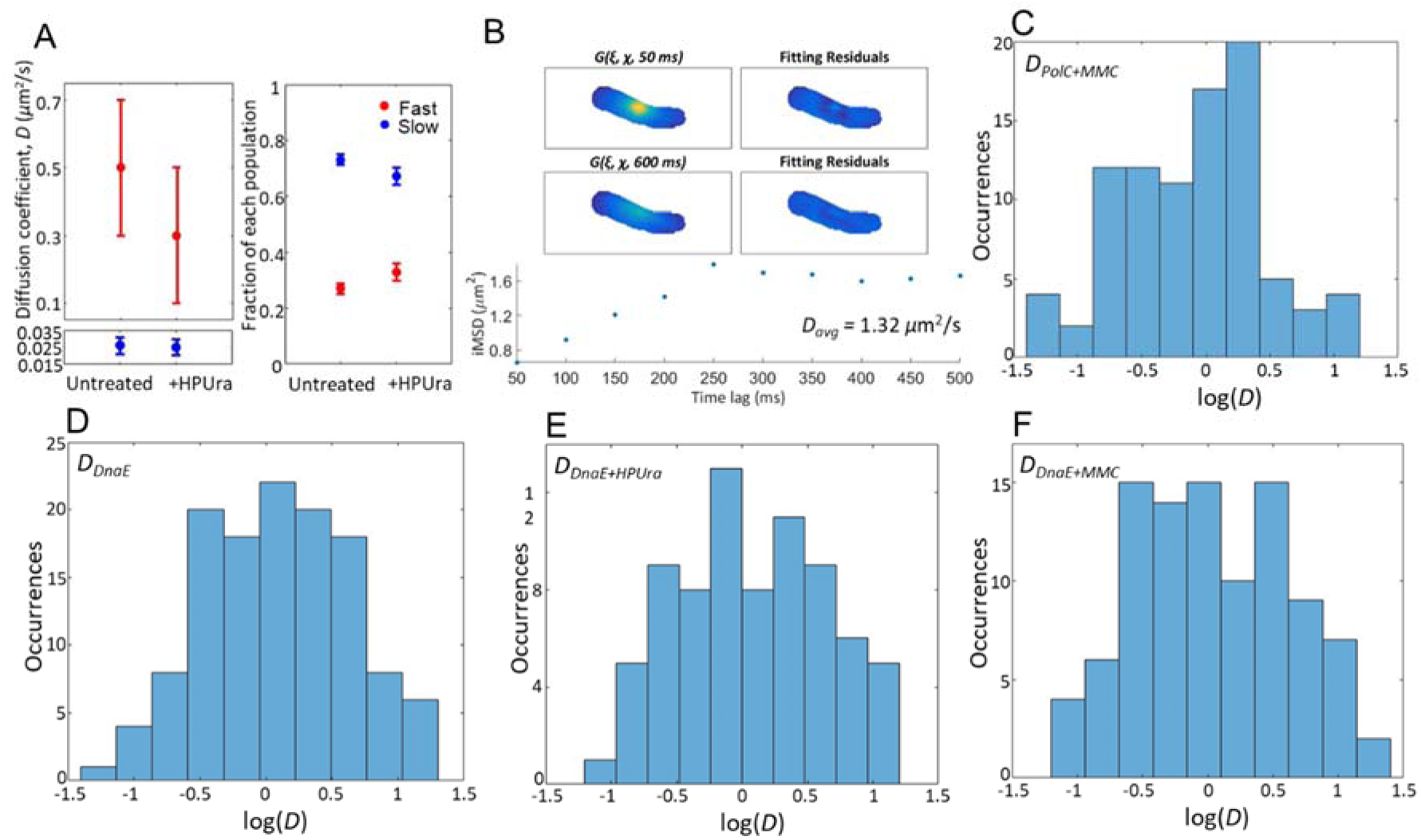
Quantification of the diffusion of PolC and DnaE in live *B. subtilis* cells. (A) Diffusion coefficients and fractional contributions from CPD analysis. Left: Diffusion coefficients, *D*_*PolC*-*fast*_ and *D*_*PolC*-*slow*_, for the ‘fast’ and ‘slow’ populations of PolC-PAmCherry in untreated and HPUra-treated cells, respectively calculated from the MSD curves in SI Fig. S1. Right: Fraction of the ‘fast’ and ‘slow’ populations in the cells under both conditions. Error bars indicate 95% confidence interval. (B) Diffusion coefficient estimation of fast-moving molecules by STICS. The STICS correlation function is computed and then fit to a Gaussian function, *G*. The long-axis variances of the Gaussian fits to the correlation functions (*iMSD*) were plotted as a function of time lag, ***τ*** (dots), and this *iMSD* curve was fit to a model for square-confined diffusion (Eq. 6) to obtain a single average diffusion coefficient measurement for each cell. (C) Distribution of the average PolC diffusion coefficient measured in each of 88 different MMC-treated cells estimated by STICS. (D) – (F) Distributions of the average DnaE diffusion coefficients measured in (D) untreated, (E) HPUra-treated, and (F) MMC-treated cells estimated by STICS.

**Table 1.** Summary of measured diffusion coefficients (in μm^2^/s) for PAmCherry-labeled proteins in untreated *B. subtilis* cells and in cells treated with HPUra or MMC. The fast and slow diffusion coefficients, *D_fast_* and *D_slow_*, respectively, are measured by single-molecule tracking and CPD analysis based on Eqs. (2) – (3). The average diffusion coefficients, *D_avg_*, are measured by STICS based on Eq. (6). For cases where two populations could be measured by CPD analysis, the population fraction is indicated in parentheses. Error bars indicate 95% confidence interval (CPD analysis) or standard error on the mean (s.e.) (STICS analysis).

|  | Untreated | +HPUra | +MMC |
| --- | --- | --- | --- |
| **PolC** | $D_{PolC-fast} = 0.5 \pm 0.2$ (27%)<br>$D_{PolC-slow} = 0.026 \pm 0.005$ (73%) | $D_{PolC-fast} = 0.3 \pm 0.2$ (33%)<br>$D_{PolC-slow} = 0.025 \pm 0.005$ (67%) | $D_{PolC,avg} = 1.4 \pm 0.2$ |
| **DnaE** | $D_{DnaE,avg} = 1.9 \pm 0.2$ | $D_{DnaE,avg} = 2.1 \pm 0.3$ | $D_{DnaE,avg} = 2.0 \pm 0.2$ |
| **DnaX** | $D_{DnaX-fast} > 1$<br>$D_{DnaX-med} = 0.2 \pm 0.1$ (33%)<br>$D_{DnaX-slow} = 0.026 \pm 0.007$ (67%) | $D_{DnaX-fast} > 1$<br>$D_{DnaX-med} = 0.17 \pm 0.02$ (45%)<br>$D_{DnaX-slow} = 0.031 \pm 0.007$ (55%) | $D_{DnaX-fast} > 1$<br>$D_{DnaX-med} = 0.16 \pm 0.06$ (36%)<br>$D_{DnaX-slow} = 0.025 \pm 0.006$ (64%) |

DnaE, which is a smaller protein than PolC, is the other polymerase essential to DNA replication in *B. subtilis* (13). We photoactivated 1 – 3 DnaE-PAmCherry molecules per cell at a time followed by imaging with microscopy. Unlike PolC, DnaE does not show measurable dynamics. Rather, although DnaE-PAmCherry molecules were indeed photoactivated (SI Fig. S2), no DnaE-PAmCherry could be tracked at our imaging rate of 50 ms/frame (Movie S2). Moreover, DnaE-PAmCherry trajectories could not even be visualized upon imaging at a faster rate of 20 ms/frame. This observation indicates extremely rapid diffusion of these DnaE polymerase molecules, which means that DnaE molecules are mobile in the cytoplasm most of the time, and the amount of time DnaE spends at the replisome is shorter than the imaging speed and far shorter than what we observe for PolC. The absence of any significant dwell time is interesting in the context of *in vitro* data showing that DnaE is responsible for extending the lagging strand RNA primers followed by a hand-off to PolC (3). Our results indicate that the DnaE to PolC hand-off occurs in less than 10 ms, or after only a few base pairs are added *in vivo*. Our striking results are most supportive of a model where DnaE extends primers during DNA replication (3). Our results do not support the model that DnaE provides a more substantial contribution to the elongation phase of chromosomal replication (19). If DnaE had such a role, we would expect the DnaE localization data to have been very similar to what we observe for PolC.

We applied spatiotemporal image correlation spectroscopy (STICS) to our DnaE-PAmCherry data to quantify the DnaE dynamics in the absence of measurable single-molecule trajectories (Fig. 2B); this analysis uses image correlation instead of tracking information to calculate the average diffusion coefficient in a cell (31,32). STICS calculated a very rapid average DnaE-PAmCherry diffusion coefficient, *D_DnaE,avg_* = 1.9 ± 0.2 μm^2^/s (Fig. 2D, Table 1).

Because DNA replication in *B. subtilis* requires the cooperation of many proteins (3), the dynamics of all replication machinery proteins, including the *β*-clamp loader component DnaX, must be coordinated at the replisome. We photoactivated 1 – 3 DnaX-PAmCherry molecules per cell at a time and visualized the molecules in our microscope (Table 1). Three types of DnaX-PAmCherry motions were observed. The fast motion is too fast to be tracked, even when imaging with a fast rate of 20 ms/frame, and STICS could not be used to quantify this fast diffusion coefficient due to the presence of many much slower molecules. Based on our frame rate, we determine a lower bound: *DDnax-fast* > 1 μm^2^/s. The medium population, representing about 33% of the trackable molecules, has a diffusion coefficient of *D*_*DnaX*-*med*_ = 0.2 ± 0.1 μm^2^/s, which is on the same order of magnitude as *D*_*PolC*-*fast*_ and which therefore must correspond to DnaX molecules transiently associating with DNA.

The slow DnaX-PAmCherry population (*D*_*DnaX*-*slow*_ = 0.026 ± 0.007 μm^2^/s) demonstrates obvious dwelling events (Movie S3), in which a single molecule remains at a sub-diffraction-limited position for a significant amount of time (> 100 ms). To understand this dwelling behavior, we quantified the dwell time of DnaX-PAmCherry. Here, consecutive steps with displacements < 100 nm are considered to demonstrate dwelling, since DnaX foci and thus the replisome exhibit only subtle motion in *B. subtilis* (radius of gyration = 84 ± 20 nm (20)). We uncovered the true dwell time with time-lapse imaging (Methods) because the PAmCherry tag photobleaches on the same seconds timescale as the DnaX dwell times. We measured an average DnaX dwell time of ***τ***_*DnaX*-*dwell*_ = 2.63 ± 0.97 s (Fig. 3A-B; Table 2), which is similar to previous reports of the PolC dwell time in *B. subtilis* (***τ***_*PolC*-*dwell*_ = 2.79 ± 0.41 s) (20). These dwell times correspond to the time needed to synthesize ~1500 DNA nucleotides, which is on the order of the length of a single Okazaki fragment synthesized on the *B. subtilis* lagging strand (3).

**Table 2.**
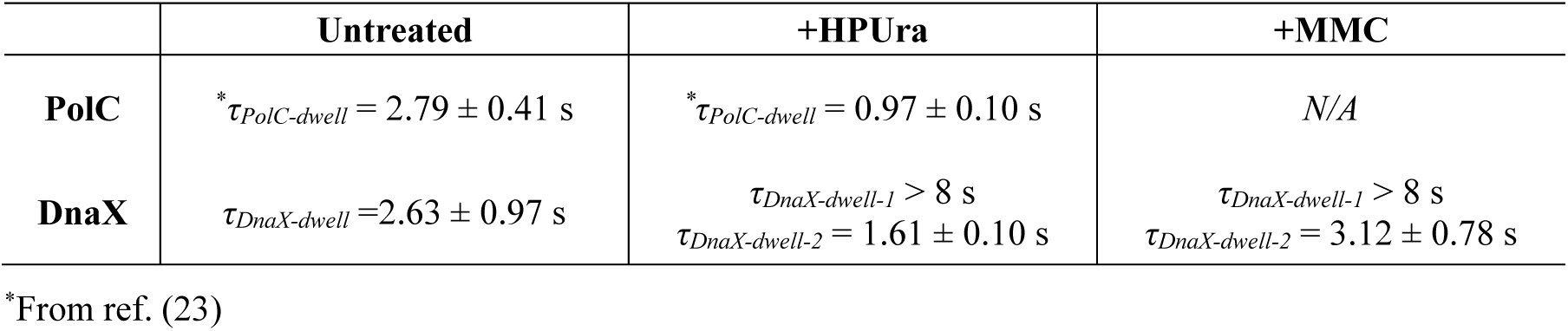
Summary of measured dwell times for PAmCherry-labeled proteins in untreated *B. subtilis* cells and in cells treated with HPUra or MMC. Error bars indicate 95% confidence interval.

**Figure 3.**
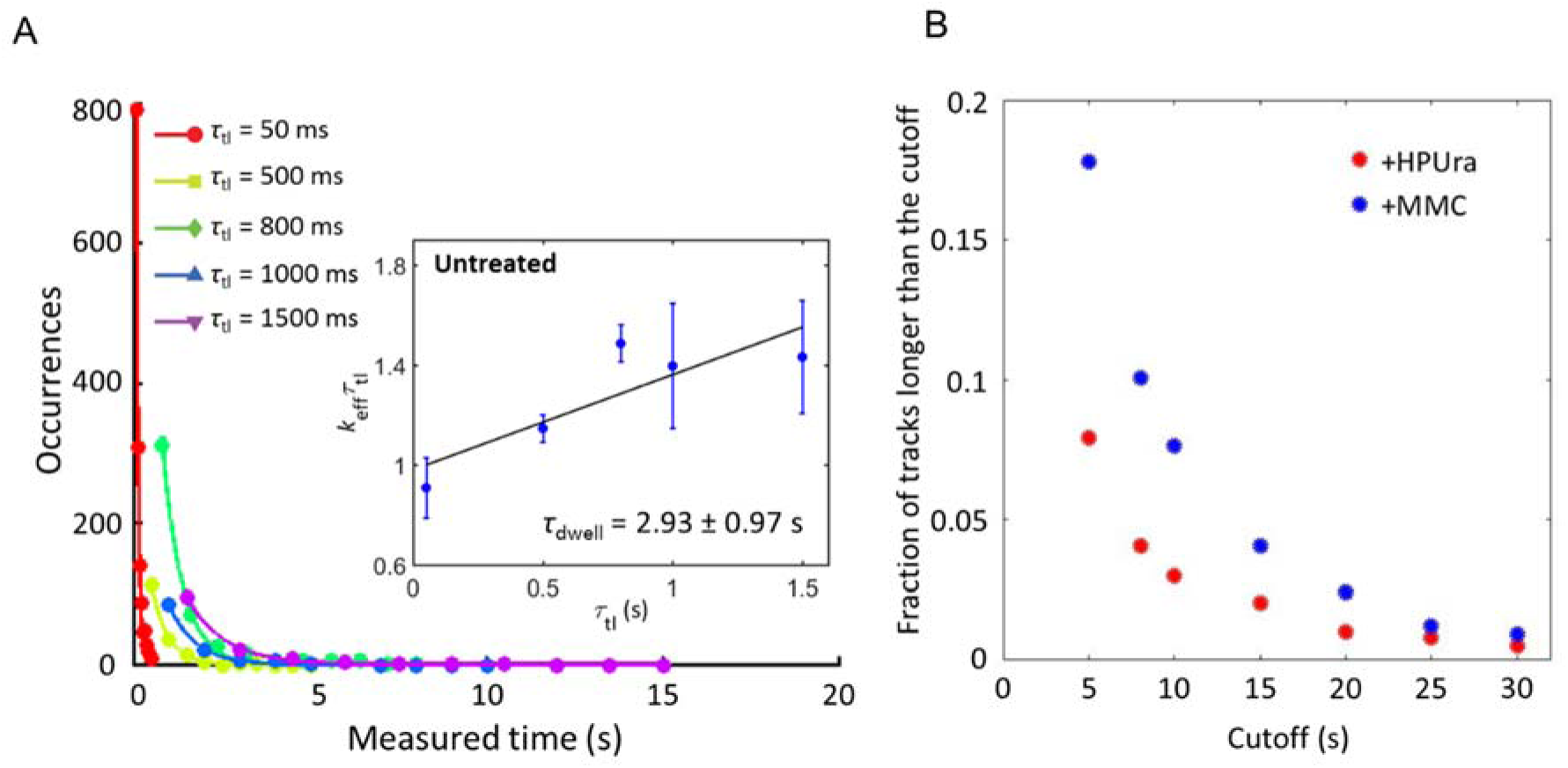
Characterization of the DnaX-PAmCherry dwelling behavior in live *B. subtilis* cells. (A) Dwell time distributions for DnaX-PAmCherry in untreated cells. The colors correspond to the time-lapse period (***τ***_*tl*_), which is the sum of the integration time and the delay time. The distributions were fit to an exponential decay function (Eq. (4)) to calculate *k_eff_*. (*Inset*) Linear fit (black line) of *k*_*eff*·***τ***_*tl*__ versus ***τ***_*tl*_, from which the dissociation rate constant, *k_off_*, the photobleaching rate constant *k_b_*, and the dwell time constant, ***τ***_*dwell*_, are obtained (Eq. (5)). Errors bars are from four rounds of bootstrapping. (B) Fraction of long tracks in HPUra-treated and MMC-treated DnaX-PAmCherry cells. Each point shows the fraction of the tracks longer than the cutoff time when **τ**_*tl*_ = 1500 ms.

### Dynamics of Replisome Proteins under Replication Arrest

To understand the mechanism that gives rise to these dynamics of motion and exchange at the *B. subtilis* replisome, we treated the cells with 6-hydroxy-phenylazo-uracil (HPUra), a drug that immediately arrests DNA synthesis by binding to the active site of PolC (33). We previously showed that HPUra treatment arrests DNA replication while maintaining the replisome complex: DnaX-mCherry foci persist after HPUra treatment (20). Here, we find that DNA replication arrest by HPUra does not arrest <u>all</u> replisome protein dynamics.

After HPUra treatment, we imaged PolC-PAmCherry, reconstructed PALM super-resolution images (Fig. 1C), and tracked PolC-PAmCherry in living *B. subtilis* cells (Fig. 1D). HPUra treatment did not significantly change the PolC dynamics: the motions are still separated into two distributions—one fast and one slow—and the CPD of the squared step sizes is fit by nearly identical diffusion coefficients and population weights (Table 1, Fig. 2A, SI Fig. S3). Taking this unchanged motion in the context of our previous reports that PolC has a shorter dwell time (faster exchange rate) after HPUra treatment (34) implies that PolC can still bind to the replisome after HPUra arrests replication, but that PolC is more likely to dissociate after HPUra arrest because it cannot add new nucleotides as synthesis is arrested. The PolC-DNA interaction is therefore stabilized by the active replication process.

Upon adding HPUra to arrest replication in *B. subtilis* cells expressing DnaE-PAmCherry, the DnaE still moved too quickly for tracking by single-molecule localization. We measured a diffusion coefficient of *D_DnaE,avg_* = 2.1 ± 0.3 μm^2^/s by STICS (Fig. 2E, Table 1). This diffusion coefficient, which is essentially unchanged from *D_DnaE,avg_* in untreated cells, indicates free diffusion of DnaE in the cytoplasm. The fact that replication arrest does not give rise to DnaE dwelling is consistent with DnaE failing to provide a major role in elongation of replication forks *in vivo*.

Upon HPUra treatment of *B. subtilis* cells expressing DnaX-PAmCherry, three mobile DnaX populations were still observed (Table 1). As in the untreated cells, the fastest population remains too fast for the diffusion coefficient to be measured, while the slowest population shows dwelling. However, after HPUra treatment, this dwelling behavior is changed (Movie S4): some DnaX molecules dwell for a very long time period (***τ***_*DnaX*-*dwell*-*1*_ > 8 s) (Fig. 3C, Table 2); the full extent of these dwells is not quantifiable with our time-lapse imaging because the typical length of a movie is 2 – 4 min and the longest practical time-lapse interval is 5 s. Sample drift or even cell elongation could occur for longer (> 5 min) movie lengths, causing the determination of dwelling events to be less certain. The average residence time for the rest of the DnaX dwelling (calculated using only tracks that dwell less than 8 s), is ***τ***_*DnaX*-*dwell*-*2*_ = 1.61 ± 0.10 s (Fig. 3D), which is shorter than the dwell time of DnaX in untreated cells and consistent with the increased PolC exchange rate after HPUra treatment (34): like most of the DnaX, PolC is more rapidly exchanged at the replisome upon replication arrest (***τ***_*PolC* -*dwell*_ = 0.97 ± 0.10 s (34)).

Unlike PolC however, the observation of two different DnaX dwell times indicates that DnaX has two different localization properties in HPUra-treated cells: after HPUra causes replication arrest, DnaX might be used in different ways to carry out different functions. We suggest that the DnaX molecules with the 1.6-s dwell times are still binding to the original sites in the replisome and exchanging faster in the same manner as PolC. This accelerated exchange rate demonstrates that the DnaX and PolC exchange dynamics are coupled at the replisome. In contrast, the longer DnaX dwell times are observed only after replication arrest and only for DnaX (not for either DNA polymerase). We therefore hypothesize that these long dwells are related to DNA repair or attempted translesion synthesis when PolC synthesis is blocked. Upon replication arrest, some DnaX molecules could be required to reload the *β*-sliding clamp for translesion DNA polymerase usage in response to HPUra.

To further investigate the effect of perturbing DNA synthesis, we treated the cells with mitomycin C (MMC), a drug that has been shown to arrest DNA replication in *B. subtilis* (35). As opposed to HPUra, which arrests replication by binding to the PolC active site (33), MMC arrests replication by inducing DNA crosslinks and monoadducts that impair fork progression (36). Contrary to HPUra-induced replication arrest, MMC treatment leads to PolC-PAmCherry molecules that are extremely fast moving: the number of single PolC-PAmCherry molecules that can be localized and fit is decreased (Fig. 1E) and the molecules are not trackable (Fig. 1F). We therefore used STICS to quantify the average diffusion coefficient of PolC, and measured to *D_PolC,avg_* = 1.4 ± 0.2 μm^2^/s (Fig. 2C, Table 1). This large average diffusion coefficient is far greater than even the fastest average diffusion coefficient for PolC-PAmCherry in the untreated cells (*D*_*PolC*-*fast*_ = 0.5 ± 0.2 μm^2^/s), which indicates that MMC treatment causes PolC-PAmCherry to dissociate from the replisome and become mobile in the cytoplasm. Furthermore, in MMC-treated cells, DnaE-PAmCherry still diffuses rapidly in the cytoplasm, with a diffusion coefficient of *D_DnaE,avg_* = 2.0 ± 0.2 μm^2^/s (Fig. 2F, Table 1). This *D_DnaE,avg_* is faster than *D_PolC,avg_* because PolC is a larger protein than DnaE. Overall, neither of these two DNA polymerases can associate with the replisome after MMC induces DNA damage in *B. subtilis* indicating that MMC damage is either very disruptive to the architecture of the replisome *in vivo* or that the repeated recruitment of translesion DNA polymerases PolY1 and PolY2 prevent association of PolC with the replisome (37,38).

Finally, DnaX-PAmCherry in MMC-treated cells still exhibited three mobile DnaX populations: *D_Dnax-fast_* was too fast to be tracked, *D*_*DnaX*-*med*_ = 0.16 ± 0.06 μm^2^/s, and *D*_*DnaX*-*slow*_ = 0.025 ± 0.006 μm^2^/s (Table 1). The exchange dynamics of DnaX-PAmCherry in MMC-treated cells were measured by single-molecule time-lapse tracking and found to be similar to the dynamics of DnaX-PAmCherry in HPUra-treated cells. As in the case of HPUra treatment, here we observed two different dwell time distributions for the slow-moving DnaX. The first dwell was too long to measure with our time-lapse imaging (***τ***_*DnaX*-*dwell*-*1*_ > 8 s). The second dwell time was calculated by using only tracks that dwell shorter than 8 s (***τ***_*DnaX*-*dwell*-*2*_ = 3.12 ± 0.78 s) (Fig. 3D, Table 2), which is similar to the DnaX dwell time in untreated cells. This correspondence between dwell times before and after MMC treatment indicates that although MMC can cause PolC to dissociate from the replisome, the DnaX binding site in the replisome still remains, and the affinity of DnaX for this binding site is not changed by MMC treatment and the ensuing MMC adduct.

## Discussion

Here, we have paired single-molecule super-resolution microscopy with complementary data analysis techniques to characterize the dynamics of three replisomal proteins, including the two essential DNA polymerases, in live *B. subtilis* cells. Our results show that all three proteins undergo dynamic exchange: they are recruited to and released from the replisome. The similar kinetics of PolC and DnaX during normal replication imply a coupling between these two proteins in the DNA replication process, and support the biochemical data for DnaX assisting in clamp loading, which would then allow PolC to associate with the *β*-sliding clamp once loaded by DnaX (39). In contrast, DnaE, which is an essential replisomal protein, undergoes an exchange too fast to measure, indicating that a single DnaE protein only extends the lagging DNA strand for only a few ms. Therefore, our localization studies of DnaE are supportive of a limited contribution for DnaE in chromosomal replication. We suggest that the fast exchange we observe is due to DnaE exclusively extending the RNA primer on the lagging strand before hand-off to PolC for the bulk of genome replication, as has been demonstrated by *in vitro* DNA replication assays (3). More recent studies suggest that DnaE might have a more significant contribution to the elongation phase of replication. If so, then we would have expected to observe a much more substantial dwell time for DnaE at the replisome. Therefore, we conclude that DnaE has a limited, but essential role in DNA synthesis.

HPUra is a specific inhibitor of PolC that only binds to the active site of and arrests DNA replication (33,40). Upon HPUra challenge, PolC and a portion of the DnaX both have a faster exchange rate. Concurrently, another population of DnaX molecules has a very long dwell time, suggesting that a new binding site with a stronger affinity to the replisome or DNA in the vicinity of the replisome is available for DnaX. One possibility is that DnaX forms a clamp-loading complex that is long-lived at a primer terminus and becomes available following HPUra treatment and failed PolC replication.

When DNA replication is arrested by the DNA damage induced with MMC, all PolC molecules dissociate from the replisome. We interpret this result to mean that the combination of crosslinks and bulky monoadducts induce exchange of PolC with the translesion DNA polymerases PolY1 and PolY2 (37,38). Some evidence suggests that DnaE is involved in DNA repair (16). If DnaE had a substantial role in DNA repair synthesis following damage with MMC, we would have expected to observe longer dwells for DnaE. Since we did not, we suggest that the contribution of DnaE to DNA repair synthesis is minor in response to MMC treatment. In contrast, we observe long dwell times for DnaX in response to MMC. One possibility is that DnaX provides an important anchoring point to maintain the integrity of the replisome when MMC adducts are encountered. Another possibility is that the MMC challenge creates 3′ termini that require loading of the *β*-sliding clamp for repair and a subset of these clamp-loading complexes are extraordinarily stable. A stable clamp-loading complex could reflect the difficulty in repairing the double-strand crosslinks that form after MMC treatment.

## Materials and Methods

### Sample preparation and single-molecule imaging

*B. subtilis* cells are all derived from PY79 and were grown at 30 °C in S7_50_ minimal medium with 1% arabinose starting from OD_600_ ~0.1. To minimize the background fluorescence and the fluorescent impurities in *B. subtilis*, minimal medium was filtered with a 0.22-μm syringe filter on the day of imaging. For PolC-PAmCherry experiments, 0.125% xylose was added to the medium to induce the ectopically expressed DnaX-mCitrine. Cells were harvested upon reaching exponential growth phase (OD_600_ ~0.55). When used, HPUra was added to the culture immediately before imaging to a final concentration of 162 μM, or MMC was added to the culture ~40 min before imaging to a final concentration of 200 ng/mL. A 2% agarose in S7_50_ pad was sandwiched between two coverslips, which had been cleaned in an oxygen plasma (Plasma Etch PE50) for 20 minutes. 1 – 2 μL of cell culture was pipetted onto the agarose pad and then used for imaging.

For single-molecule imaging, a wide-field inverted microscope was used. The fluorescence emission was collected by a 1.40-N.A. 100× oil-immersion phase-contrast objective and detected on a 512 × 512-pixel electron-multiplying charge-coupled device detector camera (Photometrics Evolve, Princeton Instruments, Acton, MA) at frame rates of 40 ms/frame for the PolC imaging and 50 ms/frame for the DnaE and DnaX imaging. PAmCherry molecules were activated by a 200-ms exposure to a 405-nm laser with a power density of ~100 W/cm^2^ (Coherent 405-100) and imaged with a 561-nm laser with a power density of ~200 W/cm^2^ (Coherent Sapphire 561-50). For two-color experiments, DnaX-mCitrine was imaged before PolC-PAmCherry with a 488-nm laser with a power density of ~7 W/cm^2^ (Coherent Sapphire 488-50).

### Super-resolution imaging and single-molecule tracking

The precise position of the single molecules were determined by fitting the point spread function (PSF) of each single molecule to a 2D Gaussian function using home-built MATLAB code (23). PALM super-resolution reconstruction images were obtained by plotting the localization of each fit convolved with a Gaussian blur with width equal to its localization uncertainty.

Single-molecule tracking was performed by maximizing the likelihood that the two subsequent localizations are the same molecule at different time points (26). A merit value, *m*, was assigned for every possible pairing:

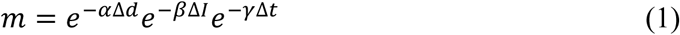

where Δ*d*, Δ*I*, and Δ*t* correspond respectively to the spatial separation, the intensity difference, and the temporal separation between the two molecules, and *α*, *β*, and *γ* are coefficients that specify how much penalty is given to each factor.

The sum of the likelihood was maximized for each set of consecutive frames, and this operation was repeated until all frames were processed. Only tracks longer than 5 frames were saved for future analysis.

### Quantification of the diffusion coefficient

To analyze the heterogeneous dynamics of PolC-PAmCherry molecules, we used the cumulative probability distribution, *P_2D_*, of PolC squared displacements, *r*^2^. Because we observe that PolC-PAmCherry can exhibit either a fast motion or a slow motion, we include the heterogeneous dynamics by fitting the distributions of *r*^2^ at each time lag, ***τ***, to a two-term model distribution (27):

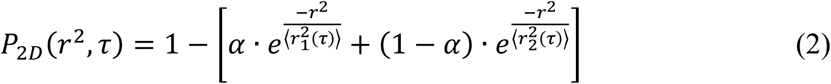

where 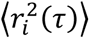 is the mean-squared displacement (MSD) for each population *i* = {1,2}, and a and (1 − *α*) account for the weight of each population.

To calculate the diffusion coefficient of each population, we fit the resulting MSD vs. ***τ*** for each of the two terms to a model that describes squared-confined motion (28,29):

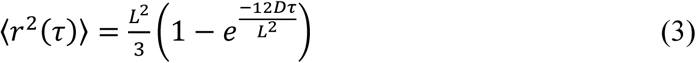

where *D* is the diffusion coefficient and *L* is the confinement length.

### Time-lapse imaging and dwell time analysis

Time-lapse imaging was used to capture the dwelling behaviors of DnaX-PAmCherry at multiple timescales. After PAmCherry was activated, every frame was captured with a 50-ms integration time, ***τ***_int_ followed by a time delay of 0 – 1.45 s (***τ***_delay_). This time lapse enables us to capture dwelling events that last up to even a few seconds despite the finite photobleaching lifetime of the fluorescent protein. The total time-lapse period, ***τ***_tl_, is defined as the sum of ***τ***_int_ and ***τ***_delay_.

The distribution of the dwell times at each ***τ***_*tl*_ were plotted (Fig. 3A) and fit to an exponential decay function, *f*

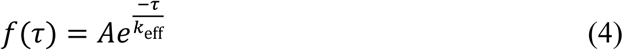

where *k*_eff_ is the effective off-rate of DnaX. Two independent factors contribute to this rate of apparent DnaX dissociation: the physical dissociation of DnaX, described by the rate constant *k*_off_, and the photobleaching of PAmCherry, described by the rate constant *k_b_* (30):

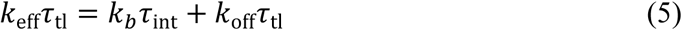

We extract *k*_off_ by fitting *k*_eff_***τ***_tl_ vs. ***τ***_tl_ to a linear function (Fig. 3B), in which the slope corresponds to the real dissociation rate constant *k*_off_. The dwell time (***τ***_well_) of DnaX is the reciprocal of *k*_off_.

### Spatiotemporal image correlation spectroscopy (STICS)

We identified the individual cell outlines with a code for cell segmentation (20), and then quantified the diffusion coefficients of molecules moving too fast to track with spatiotemporal image correlation spectroscopy (STICS) (31,32). STICS computes the correlation function of an entire fluorescence-imaging movie instead of using localization and tracking information, so it can resolve the dynamics of very fast diffusion that precludes fitting. Specifically, STICS computes the average of the spatial cross-correlations of all pairs of images separated by some time lag τ. We fit this correlation function to a Gaussian function (Fig. 2B). The long-axis variance of the Gaussian fits is called the image-mean-squared displacement (iMSD), and this quantity increases with time lag, ***τ***. The iMSD vs. ***τ*** is plotted (Fig. 2B), and fit to a model for square-confined diffusion (28):

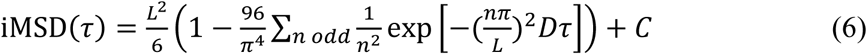

where *D* is the diffusion coefficient, *L* is the confinement length, and *C* is a constant offset. By fitting the function iMSD(***τ***) to Eq. (6) for the first 10 values of ***τ***, we obtained an average diffusion coefficient for each cell.

## Supporting Material

Three supplementary figures and four supplementary movies are available.

## Author Contributions

All authors designed the research. Y.L. and Z.C. performed single-molecule imaging experiments and data analysis. Y.L. and L.A.M. engineered the strains. All authors discussed all the results. The manuscript was written and edited by all authors. All authors read and approved the final manuscript.

## Acknowledgements

This work was supported by National Institutes of Health (NIH) grant R01-GM107312 to L.A.S. and NIH grant R21-GM128022 to J.S.B. We thank Yi Liao and Joshua Karslake for help with data analysis code. We thank Pete Burby for the codon-optimized version of PAmCherry.

## Supporting Figures

**SI Figure S1.**
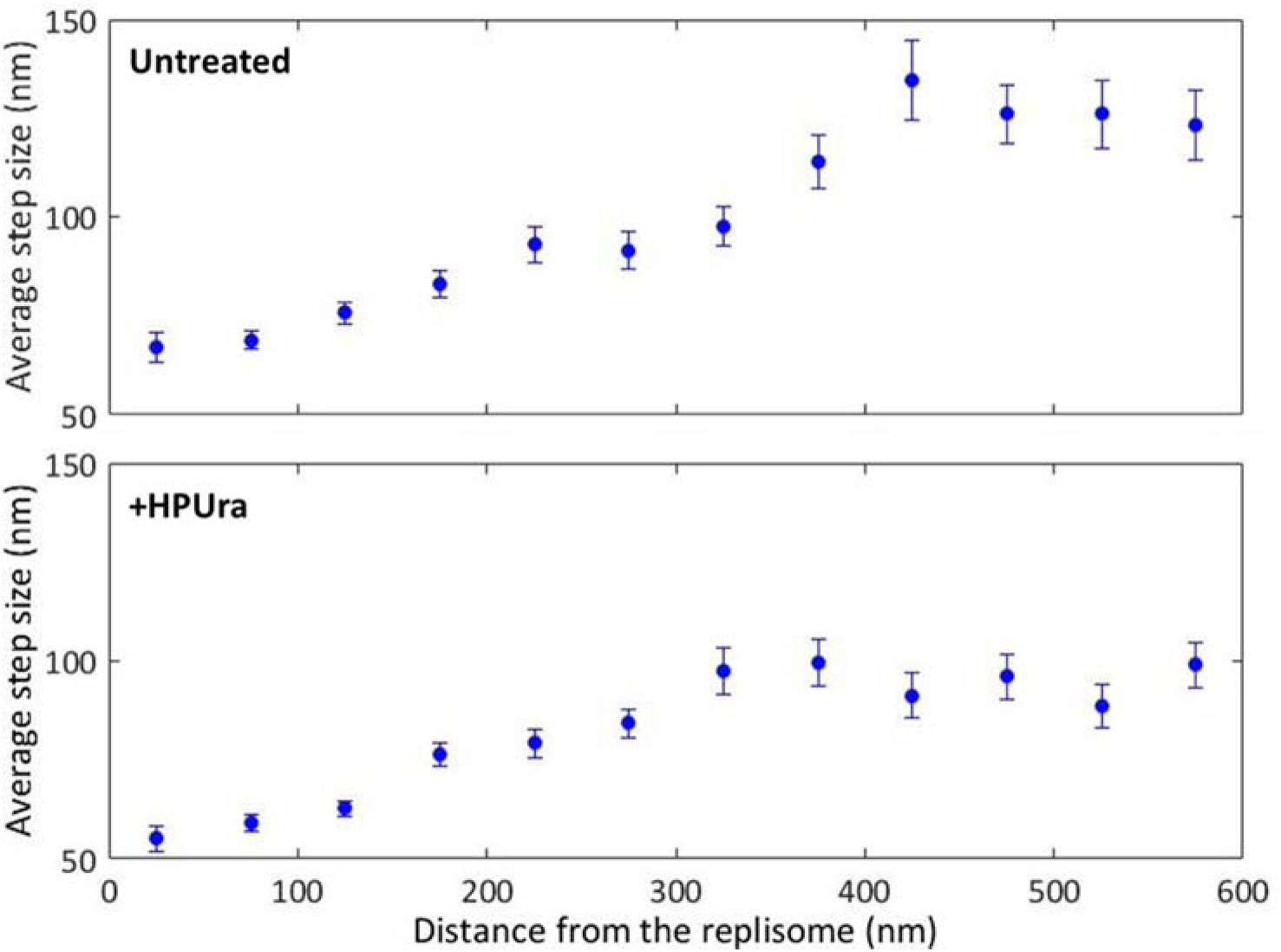
Measuring PolC dynamics as a function of subcellular positon. The mobility of PolC-PAmCherry increases as a function of separation distance from the replisome. Here, the PolC/replisome separation distance is calculated between the beginning of the PolC step to the DnaX centroid in untreated (top) and HPUra-treated (bottom) *B. subtilis* cells. Error bars: standard error on the mean (s.e.).

**SI Figure S2.**
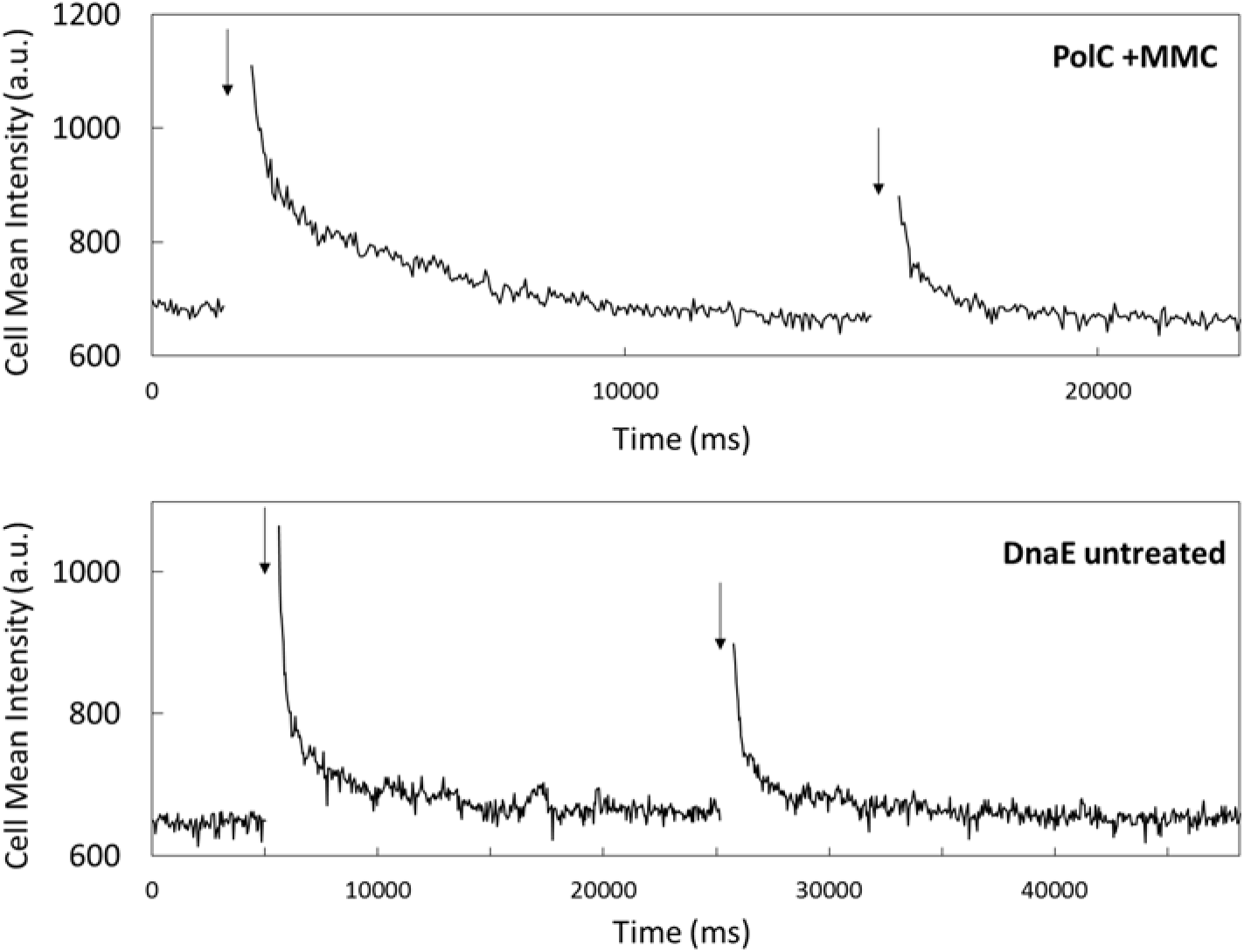
Evidence of PAmCherry photoactivation based on the time evolution of the single-cell intensity. Because PolC-PAmCherry and DnaE-PAmCherry diffusion was extremely rapid (see, e.g., SI Movie S2), very few PAmCherry single molecules were localized and no PAmCherry molecules were tracked in MMC-treated cells expressing PolC-PAmCherry or in cells expressing DnaE-PAmCherry (all conditions). Thus, we verified that PAmCherry photoactivation was in fact successful in these cells by measuring the mean cell intensity vs. time after 405-nm laser photoactivation (arrows). The increased intensity after each photoactivation in these representative data sets shows that PolC-PAmCherry (top) and DnaE-PAmCherry (bottom) were in fact successfully photoactivated. The cell intensity is calculated every 50 ms.

**SI Figure S3.**
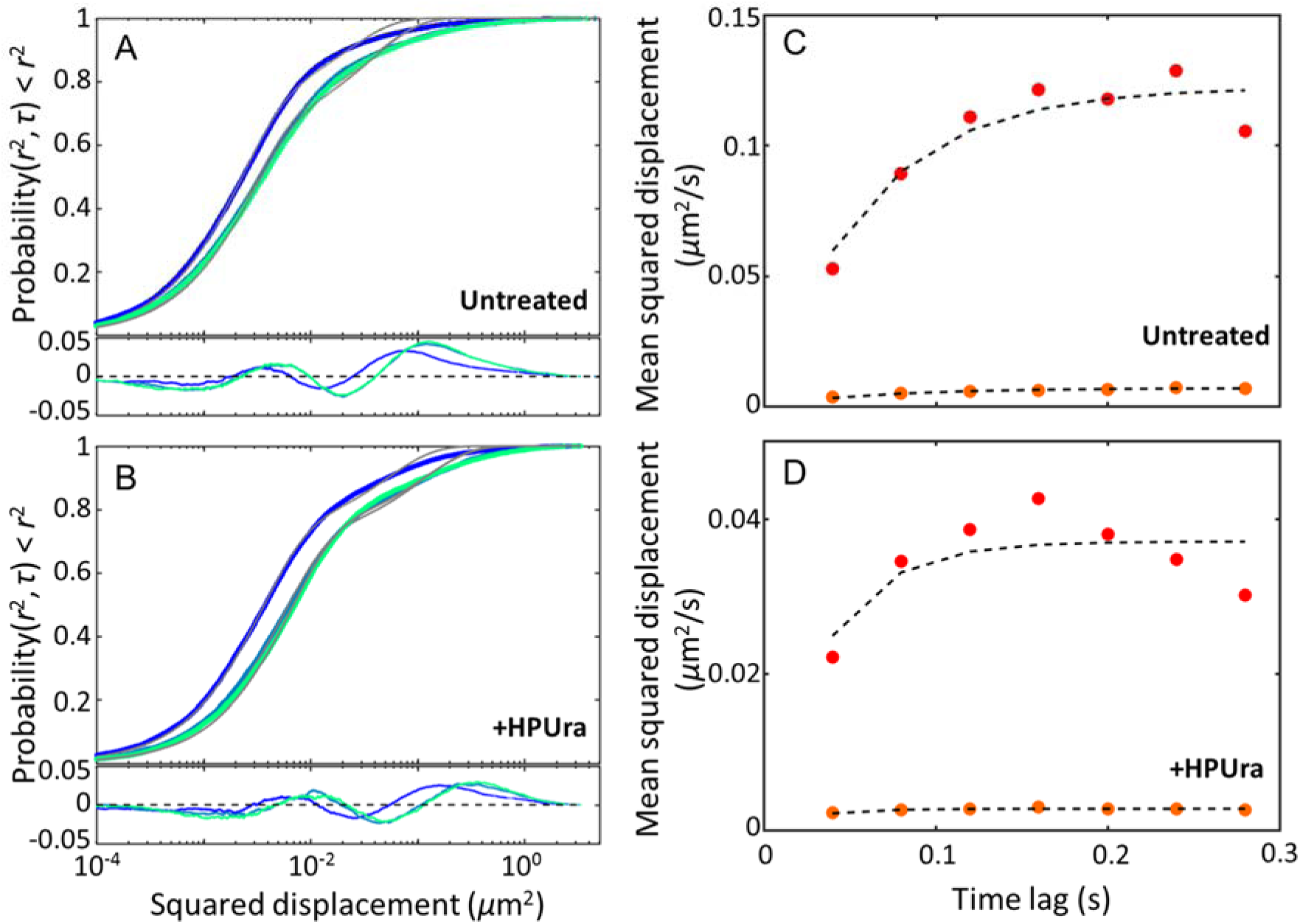
Fitting the cumulative probability distribution (CPD) of squared step sizes separates the heterogeneous PolC dynamics into the constitutive fast motion and slow motion. (A) – (B) CPD of squared displacements for each of the first three time lags, ***τ*** = {40, 80, 120 ms] for (A) PolC-PAmCherry in untreated cells and (B) PolC-PAmCherry in HPUra-treated cells. These distributions are fit to our model (Eq. 2) for two mobile, diffusive populations (grey lines) and the fit residuals are plotted below. (C) – (D) The mean squared displacement (MSD) for each population (red: ‘fast’ and orange: ‘slow’) is extracted from the fits in (A) and (B) and plotted versus time lag, ***τ***, for the first seven time lags for (C) PolC-PAmCherry in untreated cells and (D) PolC-PAmCherry in HPUra-treated cells. The MSD curves were fit to a model for square-confined diffusion (Eq. 3). The initial slopes of the curves describe the diffusion coefficients, *D*_*PolC*-*slow*_ and *D*_*PolC*-*fast*_. The saturation level of the curves is related to the confinement length, *L*, of the two populations: the slow populations are confined to *L* ~ 100 nm (similar to the replisome size (17)), whereas the fast populations are much less strongly confined (*L* ~ 500 nm).

## Supporting Movie Captions

**SI Movie S1.** PolC-PAmCherry molecules move heterogeneously in an untreated *B. subtilis* cell. Continuous imaging at 25 frames per second. Scale bar: 1 μm.

**SI Movie S2.** No DnaE-PAmCherry can be tracked in untreated *B. subtilis* cells. Continuous imaging at 20 frames per second. Scale bar: 1 μm.

**SI Movie S3.** The slow DnaX-PAmCherry population demonstrates obvious dwelling events in untreated *B. subtilis* cells. Continuous imaging at 20 frames per second. Scale bar: 1 μm.

**SI Movie S4.** After HPUra treatment, the least mobile DnaX-PAmCherry molecules dwell for > 8 s. Timelapse imaging: one 50-ms frame per 0.75 sec. Scale bar: 1 μm.

